# A 10-valent composite mRNA vaccine against both influenza and COVID-19

**DOI:** 10.1101/2024.03.05.583547

**Authors:** Yang Wang, Qinhai Ma, Man Li, Qianyi Mai, Lin Ma, Hong Zhang, Huiling Zhong, Nan Cheng, Pei Feng, Peikun Guan, Shengzhen Wu, Lu Zhang, Jun Dai, Biliang Zhang, Weiqi Pan, Zifeng Yang

**Affiliations:** State Key Laboratory of Respiratory Disease, National Clinical Research Center for Respiratory Disease, Guangzhou Institute of Respiratory Health, The First Affiliated Hospital of Guangzhou Medical University, Guangzhou, 510000, China; Guangzhou National Laboratory, Guangzhou, 510000, China; Argorna Pharmaceuticals Co., Ltd., Guangzhou, 510000, China; Respiratory Disease AI Laboratory on Epidemic and Medical Big Data Instrument Applications, Faculty of Innovation Engineering, Macau University of Science and Technology, Macau SAR, 999078, China; State Key Laboratory of Respiratory Disease, Laboratory of Computational Biomedicine, Guangzhou Institutes of Biomedicine and Health, Chinese Academy of Sciences, Guangzhou 510000, China; Guangzhou RiboBio Co., Ltd, Guangzhou 510000, China; Technology Centre, Guangzhou Customs, Guangzhou 510000, China

**Keywords:** mRNA vaccine, multi-valent, SARS-CoV-2, COVID-19, influenza

## Abstract

The COVID-19 pandemic caused by SARS-CoV-2 viruses has had a persistent and significant impact on global public health for four years. Recently, there has been a resurgence of seasonal influenza transmission worldwide. The co-circulation of SARS-CoV-2 and seasonal influenza viruses results in a dual burden on communities. Additionally, the pandemic potential of zoonotic influenza viruses, such as avian Influenza A/H5N1 and A/H7N9, remains a concern. Therefore, a combined vaccine against all these respiratory diseases is in urgent need. mRNA vaccines, with their superior efficacy, speed in development, flexibility, and cost-effectiveness, offer a promising solution for such infectious diseases and potential future pandemics. In this study, we present FLUCOV-10, a novel 10-valent mRNA vaccine created from our proven platform. This vaccine encodes hemagglutinin (HA) proteins from four seasonal influenza viruses and two avian influenza viruses with pandemic potential, as well as spike proteins from four SARS-CoV-2 variants. A two-dose immunization with the FLUCOV-10 elicited robust immune responses in mice, producing IgG antibodies, neutralizing antibodies, and antigen-specific cellular immune responses against all the vaccine-matched viruses of influenza and SARS-CoV-2. Remarkably, the FLUCOV-10 immunization provided complete protection in mouse models against both homologous and heterologous strains of influenza and SARS-CoV-2. These results highlight the potential of FLUCOV-10 as an effective vaccine candidate for the prevention of influenza and COVID-19.

**Author summary:** Amidst the ongoing and emerging respiratory viral threats, particularly the concurrent and sequential spread of SARS-CoV-2 and influenza, our research introduces FLUCOV-10. This novel mRNA-based combination vaccine, designed to counteract both influenza and COVID-19, by incorporating genes for surface glycoproteins from various influenza viruses and SARS-CoV-2 variants. This combination vaccine showed highly effective in preclinical trials, generating strong immune responses, and ensuring protection against both matching and heterologous strains of influenza and SARS-CoV-2. FLUCOV-10 represents a significant step forward in our ability to address respiratory viral threats, showcasing potential as a singular, adaptable vaccine solution for global health challenges.

## Introduction

The coronavirus 2019 disease (COVID-19) pandemic, caused by the severe acute respiratory syndrome coronavirus 2 (SARS-CoV-2), has had a substantial and multifaceted impact on global public health over the past four years. As of 8 November 2023, the World Health Organization (WHO) has documented 772 million confirmed cases and approximately 7.0 million cumulative fatalities (1). These statistics, however, are likely to be underestimated, owing to a lack of sufficient testing or poor reporting practices in the past years (2). Following a rigorous campaign involving measures such as vaccinations, medications, and restrictions on social activities, the COVID-19 pandemic has been brought under control. On 5 May 2023, the WHO lifted the status of COVID-19 from a global emergency (3); nevertheless, this declaration does not imply that the fight against infectious diseases has concluded. Numerous breakthrough infections with SARS-CoV-2 among fully vaccinated individuals suggest the potential need for annual booster vaccinations against COVID-19 (4).

Another major respiratory threat, seasonal influenza virus accounts for approximately one billion cases each year, with 290,000 to 650,000 death globally (5, 6). A notable reduction in influenza cases was observed during the 2020-2021 period, likely due to the widespread adoption of nonpharmaceutical interventions during the COVID-19 pandemic. However, the subsequent resurgence of influenza, occurring alongside SARS-CoV-2 and other respiratory diseases, has presented a dual threat to global health systems (7). This scenario is complicated further by the potential pandemic threat posed by zoonotic influenza viruses. While human infections with avian and other zoonotic influenza viruses are relatively rare, they are considerably more lethal than seasonal influenza, partly due to the absence of pre-existing immunity in the population (8, 9). For example, the highly pathogenic avian influenza A/H5N1 virus has caused 878 cases with 458 fatalities (case fatality rate: 52.2%) since its first report in 1996, and the avian influenza A/H7N9 virus has led to 1568 cases with 616 deaths (39.3%) case fatality rate: since it first emerged in 2013 (10). The resurgence of influenza, the persistence of SARS-CoV-2, and the sporadic severity of zoonotic influenza highlight the critical need for a comprehensive vaccination strategy.

In the realm of vaccine strategies, mRNA-based platforms have been distinguished by their satisfactory safety profile, high efficacy, adaptability, swift production timelines, and relatively low manufacturing costs (11, 12). Amid the COVID-19 pandemic, mRNA vaccines encoding the SARS-CoV-2 spike protein, not only received their initial authorization for human use but also rapidly became the most widely used globally, credited to their potent efficacy and expedited development timelines (13–15). However, the vaccines faced reduced effectiveness with the emergence of omicron variant (16, 17). Consequently, the mRNA vaccines were promptly adapted to include bivalent components, targeting both the ancestral and the omicron strain, and demonstrated a superior neutralizing antibody response against omicron compared to the original mRNA vaccines (18–22). The flexibility of the mRNA vaccine platform is further demonstrated by its adaptability to other respiratory diseases; for instance, mRNA-LNP vaccines encoding the HA proteins of avian influenza H10N8 and H7N9 have been shown to be highly immunogenic in phase 1 clinical trials (23), while a quadrivalent mRNA vaccine for seasonal influenza has displayed moderate to high immunogenicity in trials spanning phases 1 to 3 (24, 25).

In this study, we leveraged our established mRNA-LNP vaccine platform (11, 12) to create a novel 10-valent mRNA vaccine, aimed at targeting a diverse spectrum of respiratory pathogens. This vaccine is composed of components for all four seasonal influenza viruses (A/H1N1pdm09, A/H3N2, B/Victoria, B/Yamagata), two avian influenza viruses with pandemic potential (A/H5N1 and A/H7N9), and four strains of SARS-CoV-2 (Wuhan-Hu-1, BQ.1.1, BA.2.75.2, XBB.1.5). Subsequently, we evaluated the in vitro protein expression of this mRNA-based vaccine and confirmed its immunogenicity in mice. Moreover, we demonstrated its effectiveness in providing protection against infections from both COVID-19 and influenza.

## Results

### Design and characterization of the 10-valent mRNA vaccine (FLUCOV-10)

We developed a 10-valent mRNA vaccine candidate, named FLUCOV-10, which is designed to provide broad protection against a wide range of influenza and SARS-CoV-2 viruses (Figure 1A). The FLUCOV-10 comprises mRNAs encoding full-length HAs of each component of the quadrivalent influenza vaccines for use in the 2022-2023 influenza season in the Northern Hemisphere (i.e., A/Wisconsin/588/2019 (H1N1) pdm09, A/Darwin/6/2021 (H3N2), B/Austria/1359417/2021, and B/Phuket/3073/2013), and the HAs of two avian influenza viruses posing potential pandemic risks (i.e., A/Thailand/NBL1/2006 (H5N1) and A/Anhui/DEWH72-03/2013 (H7N9). The HA protein was selected for the influenza component due to its role as a target for neutralizing antibodies and its key function in viral entry (6). The FLUCOV-10 also includes mRNAs encoding the full-length spike proteins of ancestral SARS-CoV-2 virus and three omicron variants (i.e., BQ.1.1, BA.2.75.2, and XBB.1.5).

**Figure 1.**
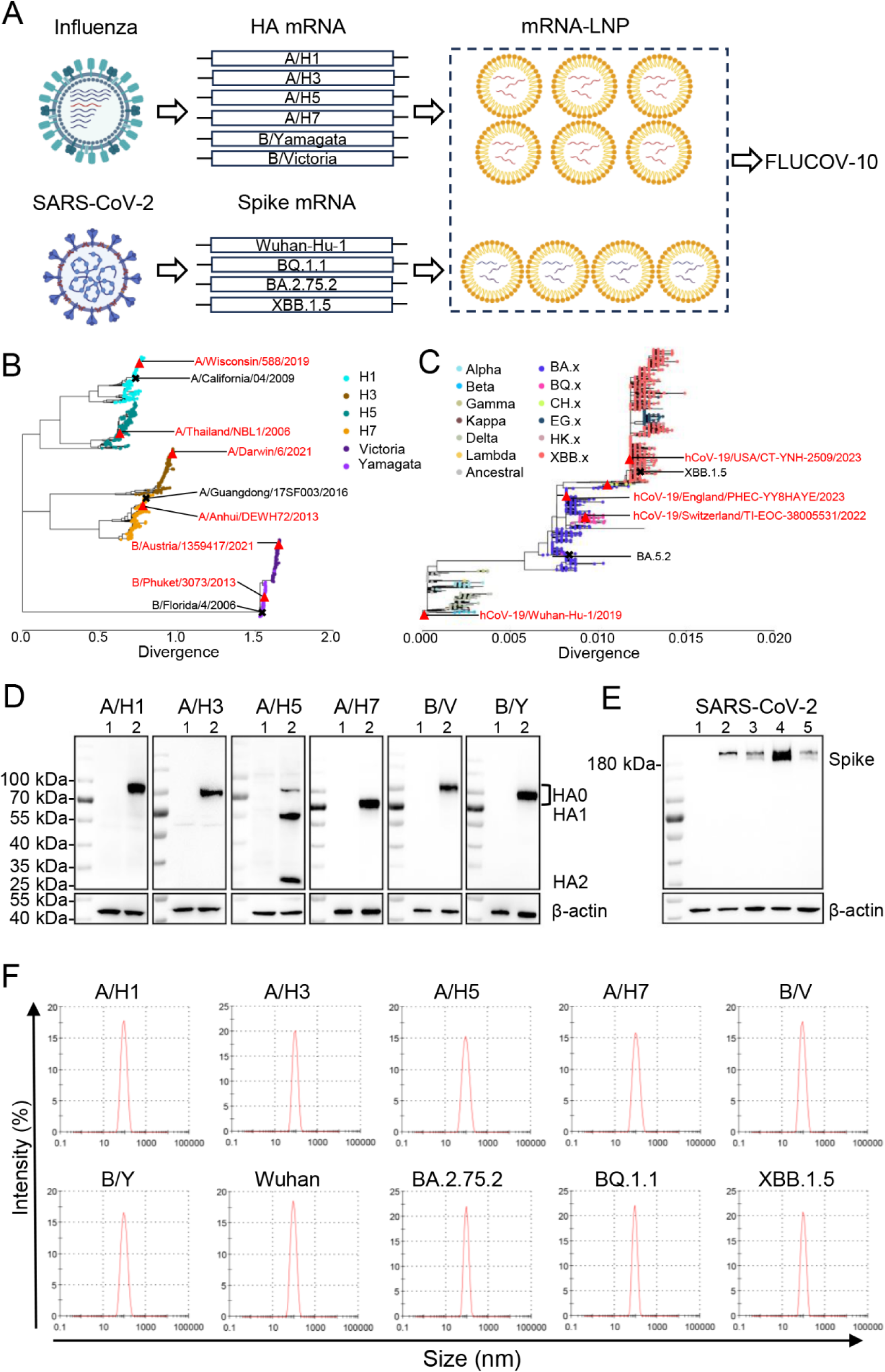
Design and characterization of 10-valent mRNA vaccine (FLUCOV-10). A. Schematic illustration of the FLUCOV-10 formulation, a 10-valent combination mRNA vaccine targeting both influenza and COVID-19. It includes mRNAs encoding the full-length HA proteins from influenza A virus subtypes A/H1, A/H3, A/H5, and A/H7, and from influenza B virus lineages B/Yamagata and B/Victoria. Additionally, it encodes full-length spike proteins from SARS-CoV-2 variants Wuhan-Hu-1, BQ.1.1, BA.2.75.2, and XBB.1.5. Each mRNA component is individually encapsulated in lipid nanoparticles (LNPs) prior to being combined into the final FLUCOV-10 formulation. B and C. Phylogenic trees were created for influenza HAs (B) and SARS-CoV-2 spikes (C) by using Nextstrain. The vaccine HAs or spike are indicated with red triangles and the challenge viruses are indicated with an “X”. D. Expression of FLUCOV-10-mRNA-encoded HA proteins in 293T cells was determined by western blotting. Lane 1, 293T cells with mock transfection; lane 2, 293T cells with indicated mRNA transfection; E. Expression of FLUCOV-10-mRNA-encoded Spike proteins. Lane 1, 293T cells with mock transfection; lane 2-5, 293T cells with Wuhan-Hu-1, BQ.1.1, BA.2.75.2, and XBB.1.5 mRNA transfection, respectively. β-actin was used as western blotting loading control.

To contextualize the sequences of vaccine strains, we conducted phylogenetic analysis using all the available HA gene sequences of influenza A/H1N1, A/H3N2, A/H5N1, A/H7N9, B/Yamgata, and B/Victoria collected since 2000, as well as all available SARS-CoV-2 spike genes (Figure 1B and C). The vaccine strains for seasonal influenza in FLUCOV-10 were selected to represent the currently circulating strains (Figure 1B), while the vaccine strains for avian influenza viruses were chosen based on the WHO’s recommendations for vaccine candidates (26). In FLUCOV-10, the inclusion of the ancestral SARS-CoV-2 strain is designed to offer cross-protection against several variants of concern, such as alpha, beta, gamma, delta and so on (11). Additionally, the incorporation of three Omicron subvariants in FLUCOV-10 was deliberately designed to address the newly emerged circulating SARS-CoV-2 variants, which possess escape properties to neutralization (27, 28) (Figure 1C).

To assess the *in vitro* expression profile of each component in FLUCOV-10, western-blotting was performed using HA- or spike-specific antibodies. As anticipated, cell lysates from the mRNA-transfected HEK293T cells exhibited a high expression level of each component (Figure 1D and E). Among the FLUCOV-10 expressing HAs, five (i.e., A/H1, A/H3, A/H7, B/Yamgata, and B/Victoria) were expressed in their precursor form (HA0), while A/H5 was present in both its precursor and cleaved forms (HA1 and HA2) (Figure 1D). This is due to the multibasic amino acid motif at the cleavage site of A/H5, which is more susceptible to cellular cleavage (29, 30). All the expressed spike proteins were maintained in their full-length form due to the intentional removal of both the furin-like cleavage motif and the S2 cleavage motif (Figure 1 E).

After encapsulating the mRNA into lipid nanoparticles (LNP), we assessed the particle size of each component in FLUCOV-10. The measurements revealed that each mRNA-LNP component consistently displayed similar average particle sizes, ranging from 90.7 to 103.9 nm (Figure 1F).

### FLUCOV-10 elicits a robust humoral immune response in BALB/c mice

To evaluate the immunogenicity of FLUCOV-10, we intramuscularly administered two doses of the vaccine to 6-8-week-old BALB/c mice, with a three-week interval between doses. Each dose contained 50 μg of mRNA, which includes 5 μg of each individual mRNA. A control group of animals was injected with a placebo. At 14 days post-booster immunization, serum samples were collected, and HA- or spike-specific antibody responses were determined by ELISA and micro-neutralization assays. The results showed that compared to placebo group, mice immunized with FLUCOV-10 produced 5,161-131,072-fold higher IgG antibody titers against all the 10 encoded HAs or spikes (*p* < 0.0001) (Figure 2A and B). In addition, the FLUCOV-10 vaccine elicited neutralizing antibodies against all the vaccine matched influenza viruses and SARS-CoV-2 viruses; whereas placebo did not induce detectable neutralizing antibodies against any of these viruses (Figure 2C and D). Intriguingly, the FLUCOV-10 induced varying levels of neutralizing antibody titers against different influenza viruses, with titers ranging from 202 to 12,902 (Figure 2C). The neutralizing antibody titers against B/Yamagata and B/Victoria were at lower levels compared to those against other influenza or SARS-CoV-2 viruses, reflecting the trend observed in their IgG titers (Figure 2A).

**Figure 2.**
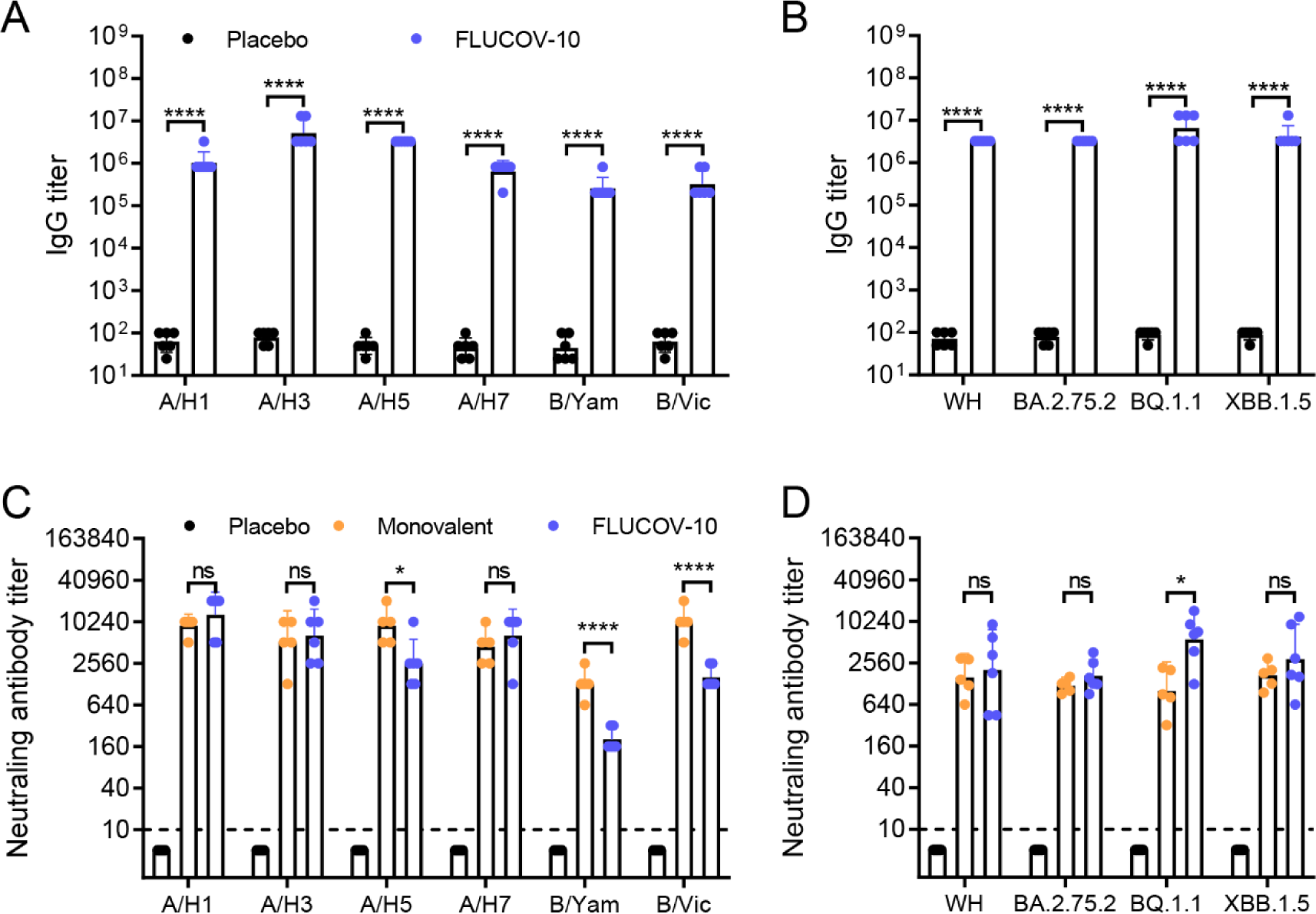
FLUCOV-10 immunization elicits a robust humoral immune response in BALB/c mice. A and B. BALB/c mice were vaccinated intramuscularly (i.m.) with the FLUCOV-10 (a combined total dose of 50 μg of mRNA, including 2.5 μg of each mRNA) or a placebo. Vaccine matched influenza HA-specific (A) or SARS-CoV-2 spike-specific (B) IgG antibody titers 14 days post the second immunization were determined by ELISA. C and D. BALB/c mice were vaccinated i.m. with FLUCOV-10, monovalent mRNA vaccines (5 μg) derived from each component of FLUCOV-10, or a placebo. Neutralizing antibody titers against vaccine matched influenza viruses (C) or SARS-CoV-2 viruses (D) were determined 14 days post second immunization by micro-neutralization assays. Data are presented as geometric means ± 95% CI (n = 5 or 6). ns, non-significant; *, *p* < 0.05; ***, *p* < 0.001; ****, *p* < 0.0001.

To explore the reason of the varied antibody responses, we simultaneously administered 5 μg doses of monovalent mRNA-LNP formulations for each component of FLUCOV-10, to BALB/c mice using the same vaccination regime. Monovalent mRNA-LNP vaccines induced neutralizing antibody titers were also at lower levels against B/Yamagata and B/Victoria compared to those against other influenza viruses (Figure 2C). Moreover, A/H5N1, B/Yamgata, and B/Victoria neutralizing antibodies were 3.5-∼6.3-fold lower in mice receiving the FLUCOV-10 vaccine compared with those receiving A/H5N1, B/Yamgata, and B/Victoria mRNA-LNPs, respectively (*p* = 0.0148, *p* < 0.0001, and *p* < 0.0001, respectively). These findings indicate that the mRNA-LNP of B/Yamaga and B/Victoria exhibited low immunogenicity and their immunogenicity was further attenuated by the presence of other components in the multivalent mRNA vaccine formulation.

In summary, two immunizations with FLUCOV-10 effectively elicited antibody responses against influenza and SARS-CoV-2 viruses.

### FLUCOV-10 elicits an antigen-specific Th1 cellular immune response in BALB/c mice

To assess the activation of HA- and spike-specific cellular immunity, we determined the antigen-specific cytokine producing splenocytes in vaccinated mice at 14 days post booster immunization by ELISpot. The results showed that the FLUCOV-10 elicited significantly higher HA- and spike-specific interferon γ (IFN-γ) and interleukin-2 (IL-2) producing splenocytes, compared to those of placebo (Figure 3A and B), while the FLUCOV-10 did not elicit higher IL-4 and IL-5 producing splenocytes (Figure 3C and D). These results indicate that the FLUCOV-10 vaccination activates Th1-biased immune responses, aligning with the observation in our previously developed mRNA vaccine platform (11, 12).

**Figure 3.**
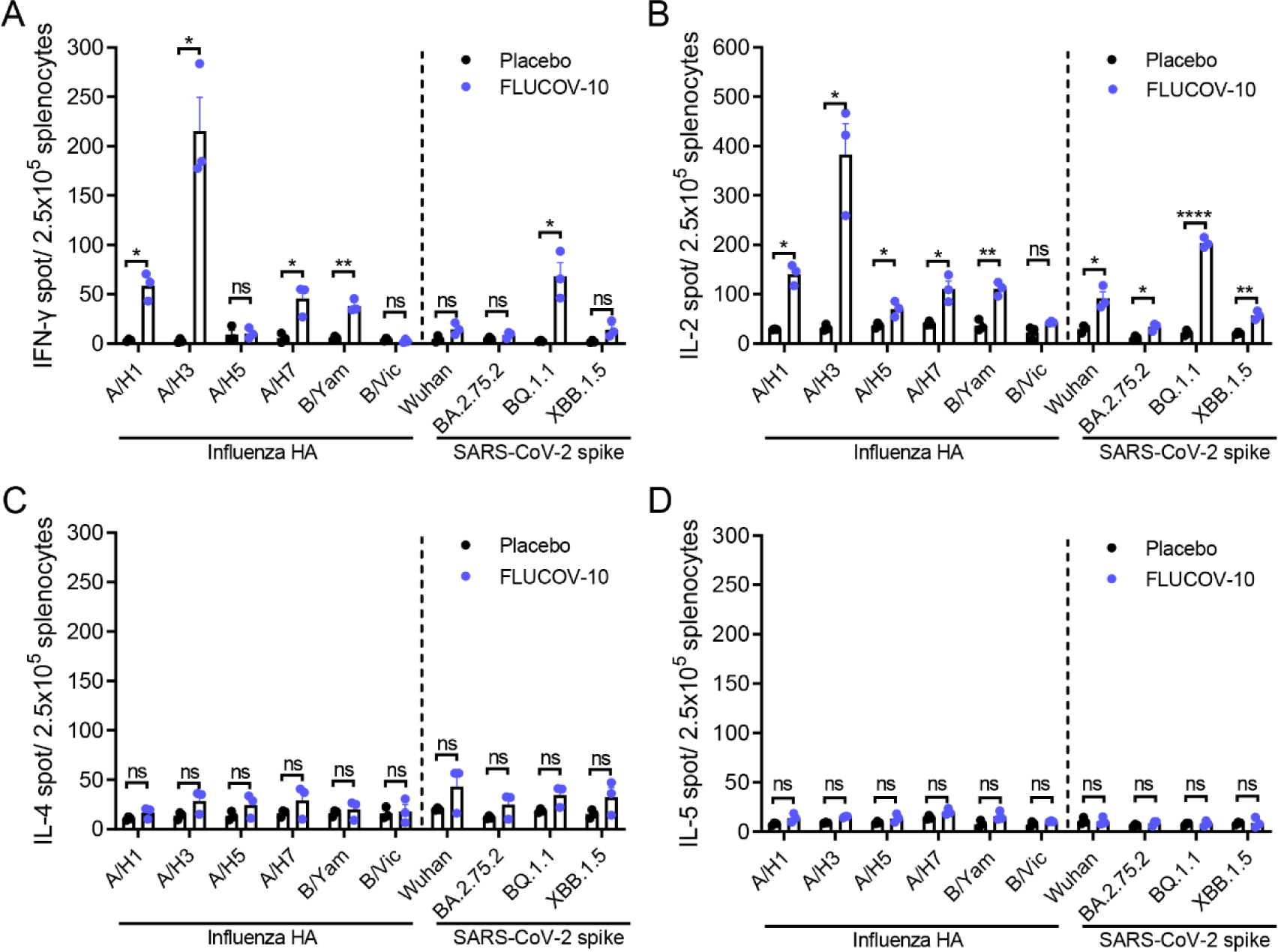
FLUCOV-10 immunization elicits an antigen-specific Th1-biased cellular immune response in BALB/c mice. BALB/c mice were vaccinated intramuscularly (i.m.) with two doses of the FLUCOV-10 or a placebo, three weeks apart. Vaccine matched HA- or spike-specific splenocytes producing IFN-γ (A), IL-2 (B), IL-4 (C), or IL-5 (D) were determined 14 days post second immunization by ELISpot. Data are presented as mean ± SEM (n = 3). ns, non-significant; *, *p* < 0.05; **, *p* < 0.01; ****, *p* < 0.0001.

Of interest, the splenocytes producing IFN-γ and IL-2 in response to FLUCOV-10 vaccination showed varying levels when stimulated with different antigens. Among the influenza HAs, A/H3-specific IFN-γ and IL-2 secreting cells reached the highest level, while those specific to B/Victoria reached the lowest level (Figure 3A and B). These results correlate with the trend observed in the subtype-specific HA IgG antibody and neutralizing antibody responses (Figure 2A and C).

### FLUCOV-10 protects mice from homologous and heterologous challenge with influenza viruses

To explore the protection efficacy against antigenically similar or heterologous influenza viruses, BALB/c mice immunized with two doses of FLUCOV-10 or placebo were challenged intranasally with A/California/04/09 (H1N1), rgA/Guangdong/17SF003/2016 (H7N9), or B/Florida/4/2006 (Yamagata lineage) three weeks after the final immunization and monitored for their weight loss and survival daily (Figure 4A). The rgA/Guangdong/17SF003/2016 (H7N9) strain was antigenically similar with the A/H7 components in FLUCOV-10 (31, 32). In contrast, the A/California/04/09 (H1N1) and B/Florida/4/2006 were both genetically and antigenically distinct from the corresponding components of FLUCOV-10, as the significant different neutralizing antibody titers were observed when comparing vaccine-matched viruses and the challenge viruses to the same mouse sera (i.e., anti-A/Victoria/2570/2019 versus anti-A/California/04/09; anti- B/Phuket/3073/2013 versus B/Florida/4/2006) (Figure 1C and Supplemental Figure 1).

**Figure 4.**
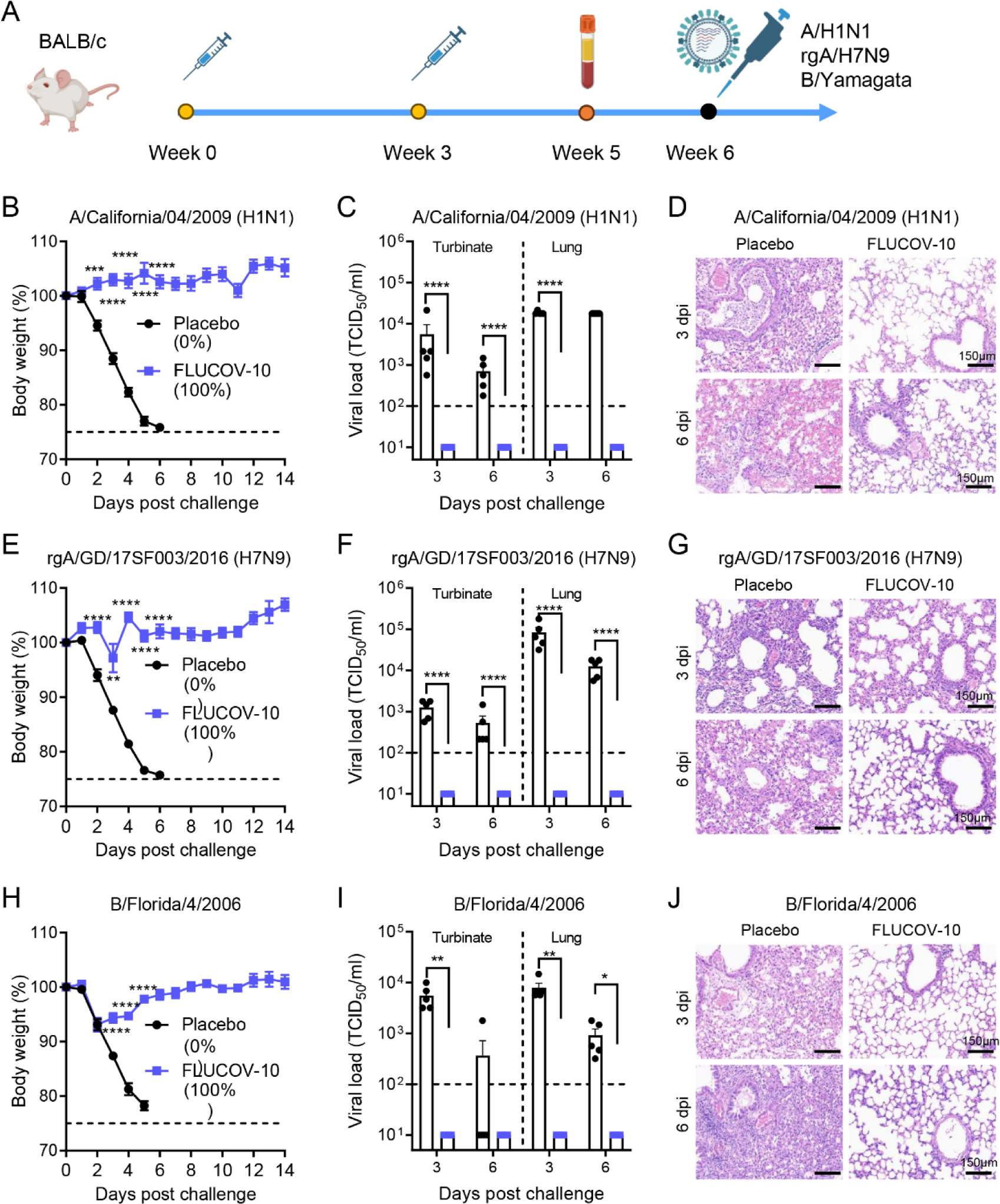
FLUCOV-10 protects mice from homologous or heterologous challenge with influenza viruses. A. Schematic diagram of the experimental design. BALB/c Mice were immunized with 50 μg of FLUCOV-10 or each volume of a placebo and boosted with the same dose after three weeks. Serum samples were collected 14 days post the second immunization. The mice were challenged 3 weeks post second immunization with 10 × mLD50 of A/California/04/2009 (H1N1) (B-D) or 10 × mLD50 of rgA/Guangdong/17SF003/2016 (H7N9) (E-G) or 3 × mLD50 of B/Florida/4/2006 (B/Yamagata) (H-J). B, E, and H, Weight changes and survival rates were recorded for 14 days (n = 7). B, E, and H, Viral titers in the turbinate or lung tissues from influenza-infected mice (n = 5 at each indicated day). C, F, and I, H&E staining of lung tissues from influenza-infected mice.

Mice immunized with the FLUCOV-10 showed significantly less weight loss than mice immunized with the placebo (*p* < 0.01) against challenge by either antigenically matched (rgA/H7N9) or heterologous virus (A/H1N1 and B/Yamagata) (Figure 4B, E, and H). Remarkably, while no survival was observed in the placebo-treated groups, the mice receiving FLUCOV-10 were completely protected against both vaccine-matched and heterologous viral challenges (Supplemental Figure 2A-C). To assess viral loads in upper and lower respiratory tract, mice were sacrificed 3 and 6 days after challenge, and lung and nasal turbinate tissues were collected for determination of viral loads by TCID50. Mice in the FLUCOV-10 groups exhibited no detectable virus in their turbinate or lung tissues at both 3 and 6 days following the challenge with either of the viruses, whereas mice from corresponding placebo groups showed significantly higher viral loads in both turbinate and lung tissues (Figure 4C, F, and I). To observe pulmonary lesions and inflammation, lung tissues at 3 and 6 days post challenge were collected for sectioning and staining. Mice from the placebo groups exhibited extensive pulmonary lesions and inflammation at both 3 and 6 days post-challenge with all three viruses (Figure 4D, G, and J). In contrast, mice immunized with FLUCOV-10 showed either mild or no apparent pulmonary lesions and inflammation following challenges with any of the viruses.

In summary, FLUCOV-10 provides complete protection against both homologous and heterologous influenza viruses, effectively preventing viral replication, lung lesions, and inflammation in the respiratory tract.

### FLUCOV-10 protects mice from challenge with SARS-CoV-2 viruses

To determine the protection efficacy against homologous and heterologous SARS-COV-2 viruses, K18-hACE2 mice immunized with two doses of FLUCOV-10 or placebo were challenged intranasally with 10^4.5^ TCID50 of XBB.1.5 and 10^4^ TCID50 of BA.5.2, respectively (Figure 5A). The XBB.1.5 strain belongs to the same clade as the vaccine component included in FLUCOV-10. In contrast, the BA.5.2 subvariant is not incorporated in the FLUCOV-10 vaccine formulation. (Figure 1C).

**Figure 5.**
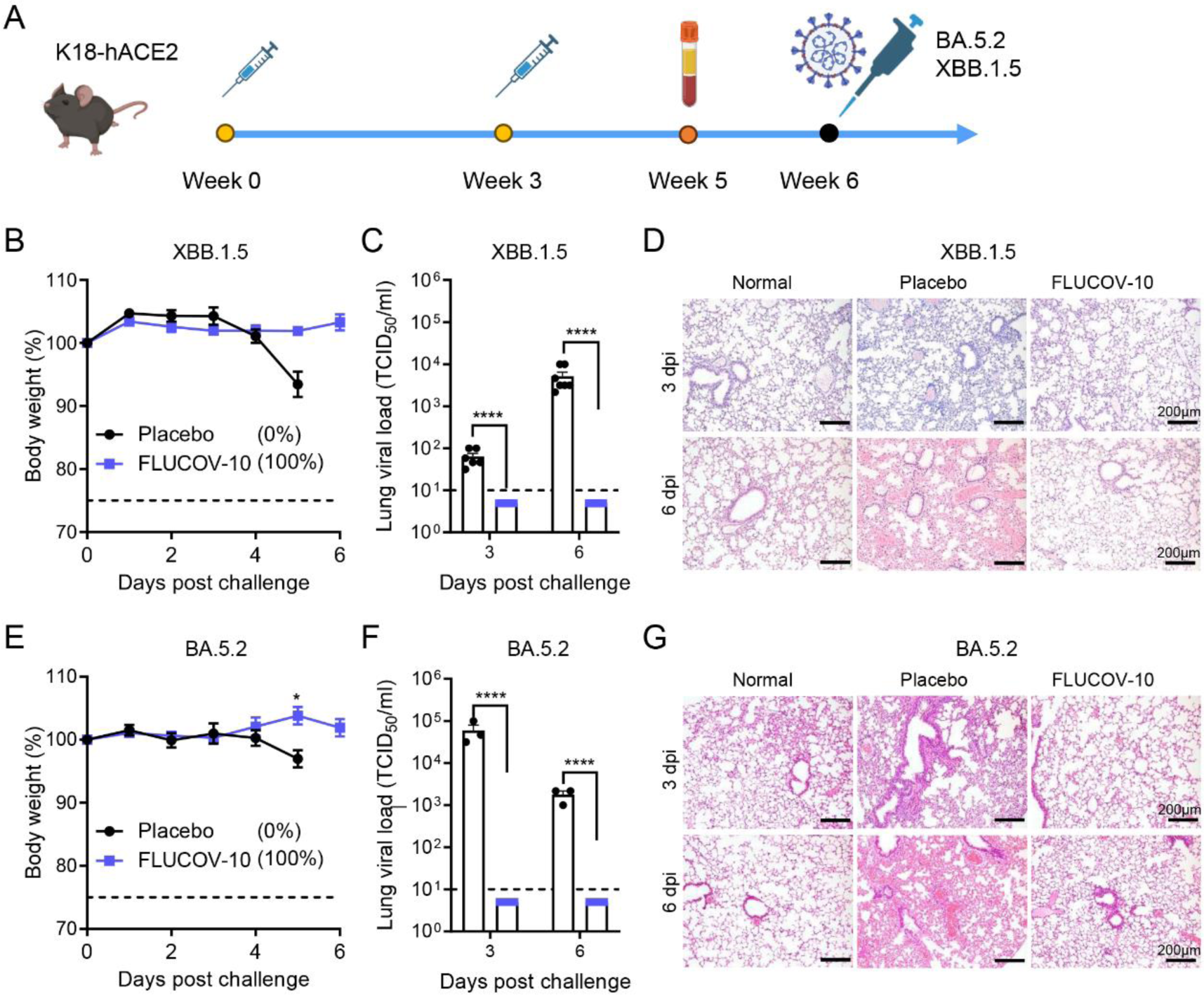
FLUCOV-10 protects mice from challenge with SARS-CoV-2 viruses. A. Schematic diagram of the experimental design. K18-hACE2 mice were immunized with 50 μg of FLUCOV-10 or each volume of a placebo and boosted with the same dose after three weeks. Serum samples were collected 14 days post the second immunization. The mice were challenged 3 weeks post the second immunization with 10^4.5^ TCID50 of hCoV-19/Chile/RM-137638/2022 (XBB.1.5) (B-D) or 10^4^ TCID50 of hCoV-19/Uganda/UG1282/2022 (BA.5.2) (E-G). B and E, Weight changes and survival rates were recorded for 14 days (n = 7 or 4). C and F: Viral titers in the lung tissues from SARS-CoV-2-infected mice (n = 5 or 3 at each indicated day). D and G, H&E staining of lung tissues from infected or normal mice.

Mice immunized with the FLUCOV-10 showed significantly less weight loss than mice immunized with the placebo against XBB.1.5 and BA.5.2 challenge (*p* = 0.0009 and *p* = 0.0128, respectively, at 5 days post challenge) (Figure 4B, E, and H). Of note, while no mice from placebo groups survived after either XBB.1.5 or BA.5.2 strain challenge, the mice receiving FLUCOV-10 were completely protected against both XBB.1.5 and BA.5.2 virus challenges (Supplemental Figure 2E and F). To assess viral loads in respiratory tract, mice were sacrificed 3 and 6 days after challenge, and lungs were collected for determination of viral loads by TCID50. Mice in the FLUCOV-10 groups exhibited no detectable virus in their lungs at both 3 and 6 days post challenge with either of the viruses, whereas mice from corresponding placebo groups showed significantly higher viral loads (Figure 4C, F, and I). Pulmonary lesions and inflammation were determined at 3 and 6 days post challenge. Mice from the placebo groups exhibited moderate to severe pulmonary lesions and inflammation at both 3 and 6 days post-challenge with both viruses (Figure 4D, G, and J). In contrast, mice immunized with FLUCOV-10 did not show apparent pulmonary lesions and inflammation following challenges with either of the viruses.

In summary, FLUCOV-10 provides complete protection against both homologous and heterologous SARS-CoV-2 viruses, effectively preventing viral replication, lung lesions, and inflammation in the respiratory tract.

## Discussion

Given the simultaneous and consecutive circulation of SARS-CoV-2 and seasonal influenza viruses, coupled with the looming threat posed by zoonotic influenza viruses, there is a pronounced and urgent need for the development of a combination vaccine targeting both SARS-CoV-2 and influenza viruses. Recently, various research groups have developed combination vaccines by using inactivated (33), recombinant protein (34, 35), and mRNA platforms (36–38). In the present study, we have utilized our previously established mRNA vaccine platform to design and assess FLUCOV-10, a universal vaccine that targets a broader range of distinct SARS-CoV-2 and influenza viruses. This vaccine comprises decavalent mRNAs encoding the full-length HAs of all four seasonal influenza viruses and two avian influenza viruses, as well as the full-length spikes of four different SARS-CoV-2 strains. This composition allows FLUCOV-10 to provide extensive protection against a wide spectrum of these respiratory viruses. To the best of our knowledge, FLUCOV-10 represents the first vaccine candidate that simultaneously targets SARS-CoV-2, seasonal, and avian influenza.

Ensuring the immunogenicity and efficacy of each component is a fundamental challenge in the development of combination vaccines (39). The FLUCOV-10 vaccine addresses this by incorporating 5 μg of each mRNA component, based on our previous reports showing that two doses of 5 μg in monovalent or bivalent mRNA vaccines achieved sterilizing immunity in mice, an effect comparable to a 20 μg dose (11, 12). We dissected the protein expression and immune response induction for each component of FLUCOV-10. Similar to previous findings (11, 12), each component in FLUCOV-10 resulted in abundant expression of HA or spike proteins in cell lysates (Figure 1 D and E), leading to robust component-specific humoral responses (Figure 2) and Th1-favored cellular responses (Figure 3) following a two-dose regimen. In response to the constantly evolving and antigenically diverse strains of influenza and SARS-CoV-2, we also evaluated the cross-reactive immunity conferred by FLUCOV-10. Our findings revealed that FLUCOV-10 produces strong neutralizing antibodies against antigenically distinct influenza viruses and inter-sublineage variants of SARS-CoV-2 (Supplemental Figure 1), surpassing known surrogate correlates of protection (40, 41). In line with expectations, animal challenge studies showed that FLUCOV-10 provided complete protection to immunized mice against both homologous and heterologous challenges of influenza and SARS-CoV-2 viruses, evidenced by significantly less body weight loss, 100% survival rates, undetectable viral loads in the respiratory tract, and absence of pulmonary lesions and inflammation. These findings suggest that FLUCOV-10 is a promising candidate vaccine, effectively targeting SARS-CoV-2, seasonal influenza, and avian influenza viruses simultaneously.

The full-length membrane-bound HA surface glycoprotein was selected for the influenza component of FLUCOV-10 due to its high immunogenicity and ability to elicit both strain-specific and cross-reactive immune responses (42, 43). Interestingly, we observed varying levels of immunogenicity among the different influenza HA components. Notably, the two mRNA components corresponding to influenza B viruses showed relatively low immunogenicity, as evidenced by their production of lower levels of IgG antibodies, neutralizing antibodies, and IFN-γ and IL-2 secreting lymphocytes, compared to those of other HA components. Observations from licensed influenza vaccines and a quadrivalent seasonal influenza mRNA vaccine candidate showed a similar pattern of lower influenza B strain responses (24, 44, 45). Upon further examination, we found that monovalent mRNA vaccines for both influenza B lineages generated significantly lower neutralizing antibodies than those for influenza A subtypes. Notably, the influenza B mRNA components in FLUCOV-10 produced even lower antibody levels compared to their monovalent counterparts. These results suggest that the reduced efficacy of influenza B components in FLUCOV-10 arises from both intrinsic factors and interactions within the other components of the vaccine. This observation warrants further investigation into optimizing the balance of components in multivalent vaccines.

The goal of a combined mRNA vaccine is to provide robust immunization across as many components as possible. Arevalo et al. reported a promising universal influenza mRNA vaccine comprising 20 HAs from 18 influenza A subtypes and 2 influenza B lineages (46). However, their study revealed that mice immunized with this 20-HA mRNA vaccine (2.5 μg per component) suffered a body weight loss of over 10% after being challenged with 5LD50 of A/California/7/2009 (H1N1pdm). In contrast, in the current study, mice vaccinated with FLUCOV-10 (5 μg per component) did not show obvious weight loss when challenged with 10LD50 of a comparable virus. This underscores the need for strategic adjustments in the number and dosage of components in combined mRNA vaccines to achieve maximum efficacy.

For the development of multivalent or combined mRNA vaccines, encapsulating each mRNA encoding separate antigens is a common approach, despite the efficiency of encapsulating all mRNAs simultaneously (36, 46–48). This method of individual LNP preparation facilitates the convenient verification of each mRNA vaccine component’s qualification, concentration, and immunogenicity (46). Additionally, in the case of combined vaccines where one component is already marketed and another is developed subsequently, it is generally necessary to manufacture each mRNA-LNP separately (47).

COVID-19 and influenza share challenges related to viral evolution and a decline in vaccine protection over time (49–52). Both diseases also exhibit seasonal trends (53), underscoring the potential need for annual booster vaccinations. Our FLUCOV-10 vaccine offers a flexible solution to these challenges, capable of swiftly adapting to emerging strains. This allows for an annual update of vaccine components to effectively combat newly emerging mutants or variants. The combination vaccine approach of FLUCOV-10 also streamlines immunization, reducing the number of injections, enhancing compliance, and minimizing adverse reactions (54, 55). This efficiency saves time for families and reduces healthcare visits, easing the burden on both individuals and healthcare systems. However, a critical aspect to consider is the phenomenon observed with repeated influenza vaccinations, where a blunted immune response and reduced vaccine effectiveness have been documented over time (56, 57). This observation highlights a critical consideration for future combination vaccine strategies involving COVID-19: the possibility that administering repeated COVID-19 vaccinations in a combination vaccine format might lead to diminishing immunogenicity. Consequently, further research is crucial to validate this hypothesis and develop and refine combination vaccine strategies to effectively tackle these dynamic and evolving viral threats.

In conclusion, our study highlights the feasibility and efficacy of a broad-spectrum mRNA vaccine, FLUCOV-10, in addressing the complex landscape of respiratory viral threats. Furthermore, the FLUCOV-10 vaccine offers a versatile and potentially effective tool in the global effort to control and prevent respiratory viral diseases. These findings underscore the value of continuing research and translation into clinical practice to establish the real-world efficacy and applicability of this vaccine approach.

## Materials and methods

### Cells

MDCK cells (CCL-34, American Type Culture Collection [ATCC]), human embryonic kidney 293T cells (CRL-3216, ATCC), and Vero E6 cells (CRL-1586, ATCC) were cultured in Dulbecco’s Modified Eagle medium (DMEM, Gibco) supplemented with 10% fetal bovine serum (FBS) at 37°C with 5% CO2.

### Viruses

Recombinant influenza viruses containing HA and NA from A/Victoria/2570/2019(H1N1)pdm09 (GISAID accession No., EPI1804971 and EPI1804972, respectively), HA and NA from A/Darwin/6/2021(H3N2) (GISAID accession No., EPI1998552 and EPI1998553, respectively), and HA and NA from B/Austria/1359417/2021(Victoria lineage) (GISAID accession No., EPI1921345, and EPI1921346, respectively) were created in the genetic background of A/Puerto Rico/8/34 (PR8) or B/Lee/40 as previously described (31). The recombinant H5N1 influenza virus containing the HA and NA genes derived from A/Vietnam/1194/2004(H5N1) and the recombinant H7N9 influenza viruses containing the HA and NA genes derived from A/Anhui/1/2013(H7N9) or A/Guangdong/17SF003/2016(H7N9) were constructed previously in our laboratory (31, 32, 58). The A/California/04/2009(H1N1) was kindly provided by Yi Shi (Chinese Academy of Sciences), the B/Florida/4/2006(Yamagata lineage) was kindly provided by Jingxian Zhao (Guangzhou Medical University), and the B/Phuket/3073/2013 (Yamagata lineage) was obtained from National Institute for Biological Standards and Control (NIBSC). All the influenza viruses were confirmed by sanger sequencing and propagated in 10-day-old embryonated eggs.

The SARS-CoV-2 viruses, including Wuhan-hu-1 (WH), BQ.1.1, BA.2.75.2, XBB.1.5, BA.5.2, were isolated from COVID-19 patients and were propagated in Vero E6 cells. All experiments involving these authentic SARS-CoV-2 strains were carried out in the BSL-3 Laboratory of the Guangzhou Customs District Technology Center.

### mRNA synthesis

The sequences encoding full-length spike proteins of SARS-CoV-2 viruses and the sequences encoding full-length HAs of influenza viruses were human codon optimized and cloned into a plasmid vector with the T7 promoter, 5’ and 3’ untranslated regions (UTRs) (59, 60), and a 120nt poly-A tail (61). To improve the spike protein’s stability and reduce protease cleavage, 2P mutations (K986P/V987P), furin cleavage site mutations (RRAR to GGSG), and S2’ cleavage site mutations (KR to AN) were introduced into its encoding sequences as described previously (11). The mRNAs were synthesized *in vitro* by T7 polymerase mediated transcription where the uridine-5′-triphosphate (UTP) was substituted with seudouridine-5’-triphosphate (pseudo-UTP). Capped mRNAs were generated by supplementing the transcription reactions with RIBO-Cap4. mRNA was purified by reversed-phase high-performance liquid chromatography (RP-HPLC) (62). RNA quality was analyzed by bioanalyzer analysis (Agilent 2200 Tape station). mRNA concentrations were measured by UV spectroscopy.

DNA sequences synthesized for this study originated from the following list of spike proteins and HAs that were included: Wuhan-Hu-1 (GenBank accession No. 43740568), hCoV-19/Switzerland/TI-EOC-38005531/2022 (BQ.1.1, GenBank accession No. OX366792.2), hCoV-19/England/PHEC-YY8HAYE/2023 (BA.2.75.2, GISAID accession No. EPI_ISL_16679125), hCoV-19/USA/CT-YNH-2509/2023 (XBB.1.5, GISAID accession No. EPI_ISL_16570202), A/Wisconsin/588/2019 (H1N1)pdm09 (GISAID accession No. EPI_ISL_404460), A/Darwin/6/2021(H3N2) (GISAID accession No. EPI_ISL_2233238), A/Thailand/NBL1/2006 (H5N1) (GenBank accession No. KJ907470:1-1707), A/Anhui/DEWH72-03/2013 (H7N9) (GenBank accession No. CY181529:22-1704), B/Austria/1359417/2021 (B/Victoria lineage) (GISAID accession No. EPI_ISL_2378894), B/Phuket/3073/2013 (B/Yamagata lineage) (GISAID accession No. EPI_ISL_161843).

### mRNA-LNP preparations

The FLUCOV-10 vaccine comprises a total of 50 μg of mRNA, distributed equally among 10 different mRNAs (5 μg each). These mRNAs encode the HA and spike antigens from six influenza viruses and four SARS-CoV-2 viruses, as detailed above. Each mRNA is separately formulated into Lipid nanoparticles (LNPs) and then mixed, prior to vialing so 10 different mRNA formulations are present in the vial. LNPs were prepared by microfluidic mixing using the previously described method (63). Briefly, lipids were dissolved in ethanol at molar ratios of 45:16:15:1.0 (Ionizable lipid: Cholesterol: DSPC: DMG-PEG2000). The lipid mixture was rapidly combined with a buffer of 50 mM sodium citrate (pH 4.0) containing mRNA at a volume ratio of aqueous: ethanol using a microfluidic mixer (PNI Nanosystems, Vancouver, BC, Canada). Formulations were dialyzed against PBS (pH 7.2) in the dialysis cassettes (Thermo Scientific, Rockford, IL, USA) for at least 18 h. Formulations were diluted with PBS (pH 7.2) to reach a required concentration, and then passed through a 0.22-mm filter and stored at 4°C until use. Formulations were analyzed for particle size by using a ZETASIZER, and the mRNA encapsulation, residues, endotoxin and bioburdens were also confirmed.

### mRNA transfection

293T cells were seeded in 12-well plates at 1 × 10^6^ cells per well and cultured at 37 °C in 5% CO2 for 16 h. 10 μg of each mRNA encoding HA or spike protein was transfected into 293T cells using riboFECTTMmRNAtransfection Reagent (C11055, Ribobio, Guangzhou, China). Cell lysates were harvested by RIPA lysis buffer (R0030, Solarbio, Beijing, China) at 48 h after transfection, and mixed with 5 × SDS-loading, following bycentrifugation at 12,000 × g. The samples were loaded for SDS–PAGE. The HA and spike proteins in cell lysates were then detected by western blotting using a mouse monoclonal antibody against SARS-CoV-2 spike proteins (GTX632604, GeneTex), a rabbit polyclonal antibody against influenza A/H1 HA (11055-T62, Sino Biological), a mouse monoclonal antibody against influenza A/H3 HA (11056-MM03, Sino Biological), a rabbit polyclonal antibody against influenza A/H5 HA (11062-T62, Sino Biological) ], a rabbit polyclonal antibody against influenza A/H7 HA (40103-T62, Sino Biological) a rabbit polyclonal antibody against influenza B/Yamgata lineage HA (11053-T62, Sino Biological), and a mouse monoclonal antibody against Influenza B/Victoria lineage HA (11053-MM06, Sino Biological). β-actin was detected using anti-β-actin antibody.

### Animal experiments

BALB/c mouse experiments were performed in accord with Regulations of Guangdong Province on the Administration of Laboratory Animals and Institutional Animal Care and Use Committee of Guangzhou Medical University (IACUC Approval No. IACUC-2023-001). Six- to eight-week-old female BALB/c mice (Guangdong Vital River Laboratory Animal Technology, Guangzhou, China) were immunized intramuscularly with 5 μg of each monovalent mRNA-LNP (in a 50 μl volume), 50 μg of FLUCOV-10 (in a 50 μl volume) or an equal volume of placebo and boosted with an equal dose at 21 days post-initial immunization. Serum samples were collected prior to initial immunization and 14 days after booster immunization. For influenza virus challenges, vaccinated mice were anesthetized and infected intranasally with 10LD50 of A/California/07/2009 (H1N1), 10LD50 of recombinant A/Guangdong/17SF003/2016 (H7N9) (referred to as rgA/Guangdong/17SF003/2016 H7N9)), or 3 LD50 of B/Florida/4/2006 in 50 μl of PBS at 3 weeks after booster immunization. Weight loss and survival were monitored for 14 days after challenge. Animals that lost more than 25% of their initial body weight were humanely anesthetized. At 3 and 6 days post challenge, mouse lungs and nasal turbinates were collected for viral titration and histological analyses.

For SARS-CoV-2 challenges, six- to nine-week-old female K18-hACE2 mice (Gempharmatech, Nanjing, China) were immunized with the same regimen as that of BALB/c mouse experiments. Three weeks post booster immunization, the mice were infected intranasally with 10^4^ TCID50 of hCoV-19/Uganda/UG1282/2022 (BA.5.2) or 10^4.5^ TCID50 of hCoV-19/Chile/RM-137638/2022 (XBB.1.5). Weight loss and survival were monitored for 6 days after challenge. At 3 and 6 days post challenge, mouse lungs were collected for viral titration and histological analyses. All work with live SARS-CoV-2 virus was performed in the Biosafety Level-3 (BSL-3) containment laboratories.

### Enzyme-linked immunosorbent assay (ELISA)

SARS-CoV-2 spike- and influenza HA-specific IgG antibody titers were determined by ELISA. 96-well plates (JET BIOFIL) were coated with recombinant spike proteins of Wuhan-hu-1 (ACROBiosystems, SPN-C52H4), BA.2.75.2 (ACROBiosystems, SPN-C522r), BQ.1.1 (ACROBiosystems, SPN-C522), or XBB.1.5 variant (ACROBiosystems, SPN-C524i), or recombinant Influenza HA proteins of A/Wisconsin/588/2019/A/Victoria/2570/2019 (H1N1) (Sinobiological, 40787-V08H1), A/Darwin/6/2021 (H3N2) (Sinobiological, 40868-V08H), B/PHUKET/3073/2013 (Sinobiological, 40498-V08B), B/Austria/1359417/2021 (Sinobiological, 40862-V08H), A/Vietnam/1194/2004 (H5N1) (Sinobiological, 11062-V08H1), or A/Hangzhou/3/2013 (H7N9) (Sinobiological, 40123-V08B) with a concentration of 2 μg/ml at 4°C overnight. The plates were washed three times with PBS containing 0.1% Tween 20 (PBST) and subsequently blocked with 1% bovine serum albumin in PBST at 37°C for 1 h. After blocking, 100 μl of serial dilutions of heat-inactivated serum sample was added to the plates, followed by incubation at 37°C for 1 h. Following thorough washes, HRP-conjugated Goat anti-mouse IgG (H+L) antibody (Proteintech, SA00001-1) was added to the plates and incubated at 37°C for 1 h. After three additional washes, 100 μL of TMB peroxidase substrate (TIANGEN, PA107-02) was added to each well and incubated for 15 min before being stopped by adding 2 M H2SO4, and the absorbance was measured at 450 nm using a TECAN Infinite M200 Pro plate reader. Endpoint titers were determined as the reciprocal of the highest serum dilution that exceeded the cut-off values (calculated as the mean ± SD of negative controls at the lowest dilution).

### Micro-neutralization (MN) assay

To determine neutralizing antibody titers against influenza viruses, mouse serum samples were treated with receptor-destroying enzyme II (RDE II) (Denka-Seiken) for 16 h at 37°C, followed by heat-inactivation for 30 min at 56°C. The MN assays were performed as previously described (31). To determine neutralizing antibody titers against SARS-CoV-2 viruses, serum samples collected from immunized mice were inactivated at 56 °C for 30 min and the MN assays were performed as described elsewhere (12). The MN titer was defined as the reciprocal of the highest serum dilution capable of neutralizing 50% of viral infections in MDCK cells (for influenza viral titers) or Vero E6 cells (for SARS-CoV-2 viral titers). The minimum MN titer detected in this study was 10; thus, for statistical purposes, all samples from which the MN titer was not detected were given a numeric value of 5, which represents the undetectable level of MN titer.

### ELISpot

Cellular immune responses were determined by using IFN-γ (Dakewe Biotech, 2210005), IL-2 (Mabtech, 3441-4HPW-2), IL-4 (Dakewe Biotech, 2210402), and IL-5 (Mabtech, 3391-4HPW-2) precoated ELISpot kits according to the manufacturer’s instructions. Briefly, Spleen lymphocytes isolated from BALB/c mice 14 days after the booster vaccination and plated at 2.5 ×10^5^ cells/well were added to the pre-coated plates. The spleen lymphocytes were stimulated with 1 μg/ml recombinant spike proteins or HA proteins and cultured at and 37°C and 5% CO2 for 20 h. Concanavalin A (Sigma) was used as a positive control, and RPMI 1640 medium (Gibco, Thermo Fisher Scientific) was used as a negative control. The plates were then washed 6 times with wash buffer and incubated for 1 h with biotinylated anti-mouse IFN-γ, IL-2, IL-4, or IL-5 antibody. Streptavidin-HRP was added to the plates and incubated for 1 h. After the final washes, the AEC substrate solution was added and stopped with water. The air-dried plates were read by using ELIspotreader.

### Infectious viral titration by TCID50

The right lung lobes were homogenized in 0.5 DMEM containing 0.3% BSA (Sigma-Aldrich) and 1% penicillin/streptomycin (Gibco, Thermo Fisher Scientific) for 1 min at 6,000 rpm by using a homogenizer (Servicebio). The turbinate was homogenized in 1 ml of the same medium. The debris were pelleted by centrifugation for 10 min at 12,000 × *g*. Their infectious virus titers were determined by TCID50 with MDCK cells (for influenza viral titers) or Vero E6 cells (for SARS-CoV-2 viral titers) as previously described (64, 65).

### Histopathology

Mouse left lung lobes were fixed in 10% buffered formalin, paraffin-embedded, sectioned at 4 µm, and stained with hematoxylin and eosin (H&E) for histopathological examination.

### Phylogenetic analysis

The phylogenetic analysis was performed as described previously (46). All available full-length HA genes collected during January 1, 2000, to June 30, 2023 for the influenza A(H1N1) (2009-present), A/H3N2 (2000-present), A/H5, A/H7, B/Yamagata, B/Victoria viruses, and spike genes for SARS-CoV-2 ancestral strains and omicron variants were downloaded from GISAID. To contextualize the sequences of vaccine strains and challenge strains, we utilized the Nextstrain pipeline (66) to build two separate phylogenetic trees: one for influenza HAs (A/H1, A/H3, A/H5, A/H7, B/Yamagata, and B/Victoria) and the other for SARS-CoV-2 spikes (30). For influenza tree, we randomly subsampled 10 sequences per HA type (for influenza A sequences) or lineage (for influenza B sequences) for each year. We excluded duplicate sequences, any sequences sampled before 2000, and sequences with incomplete collection dates or non-nucleotide characters. For the SARS-CoV-2 tree, we randomly subsampled approximately 1000 sequences from the omicron lineages based on the pre-analysis results from the Nextstrain pipeline. Following subsampling, sequences were aligned using MAFFT (67), and divergence phylogenies were constructed with IQ-TREE under a General Time Reversible (GTR) substitution model (68). Finally, tree plotting and visualization were carried out using ggtree (https://guangchuangyu.github.io/software/ggtree/).

### Statistical analyses

Statistical analyses were conducted using GraphPad Prism version 9. Data are presented as geometric means ± 95% CI for antibody titers, and means ± SEM for all other data. For statistical significance testing, an unpaired *t* test was applied when data showed equal variation between groups, and Welch’s *t* test was used for data with unequal variation. For comparisons involving multiple groups, one-way ANOVA with Tukey’s post-hoc test was employed. To achieve normality, antibody titer data were log-transformed prior to analysis. A *P* value of less than 0.05 was considered statistically significant.

## Acknowledgements

The authors thank Yi Fang for her assistance in phylogenetic analyses and thank Qiuwen Tang for his support in the animal studies. We sincerely thank Dr. George Dacai Liu for his comments and assistance in revising our manuscript. This work was supported by National Key R&D Program of China (2021YFC1712904), National Natural Science Foundation of China (82174053, 82361168672, 81761128014 and 31970884), Natural Science Foundation of Guangdong Province (2022A1515010301), Macao Science and Technology Development Fund (0022/2021/A1 and 005/2022/ALC), Guangzhou Science and Technology Bureau project (202201020523, 202201020449), The Young Top Talent of Science and Technology Innovation Department of Guangdong Province (2021TQ060189), The Youth Lift Project of China Association for Science and Technology (2020-2022QNRC001), Open Project of State Key Laboratory of Respiratory Disease (SKLRD-OP-202001 and SKLRD-OP-202209), Self-supporting Program of Guangzhou Laboratory, Grant No. SRPG22-007, and Guangdong-Hong Kong-Macao Joint Laboratory of Respiratory Infectious Diseases Funding Project (GHMJLRID-Z-202105).

## Supplementary Information

**Supplemental Figure 1.**
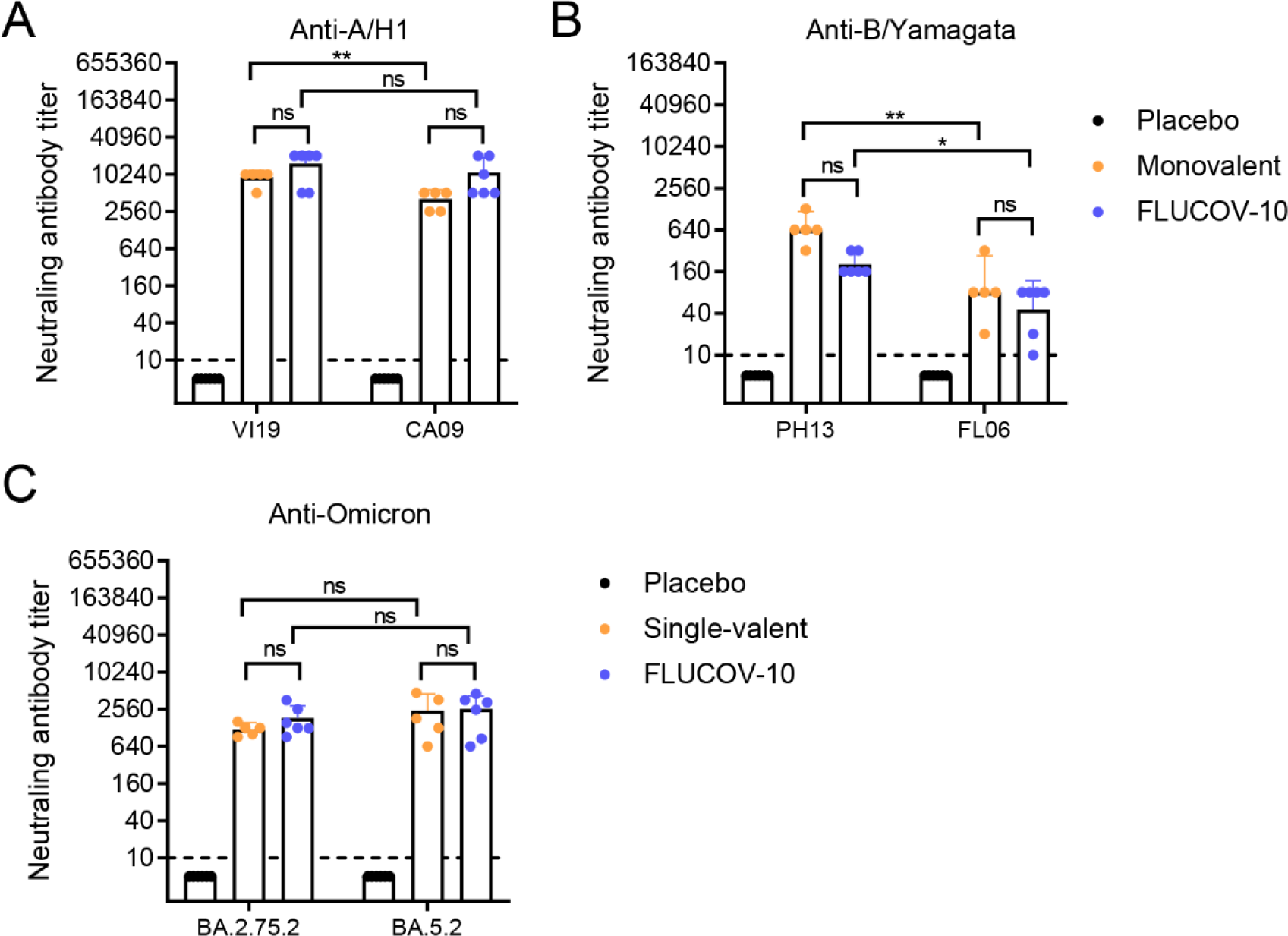
FLUCOV-10 immunization elicits a cross-reactive humoral immune response in BALB/c mice (related to Figure 2). A. BALB/c mice were vaccinated i.m. with the FLUCOV-10, monovalent A/H1 mRNA vaccines derived from FLUCOV-10 or a placebo. Neutralizing antibody titers against vaccine matched (A/Victoria/2570/2019, VI19) and antigenically distinct (A/California/04/2009, CA09) A/H1N1pdm09 influenza viruses were determined 14 days post second immunization by micro-neutralization assays. B. BALB/c mice were vaccinated i.m. with the FLUCOV-10, monovalent B/Yamagata mRNA vaccines derived from FLUCOV-10 or a placebo. Neutralizing antibody titers against vaccine matched (B/Phuket/3073/2013, PH13) and antigenically distinct (B/Florida/4/2006, FL06) B/Yamagata influenza viruses were determined 14 days post second immunization by micro-neutralization assays. Data are presented as geometric means ± 95% CI (n = 5 or 6). C. BALB/c mice were vaccinated i.m. with the FLUCOV-10, monovalent BA.2.75.2 mRNA vaccines derived from FLUCOV-10 or a placebo. Neutralizing antibody titers against vaccine matched (BA.2.75.2) and antigenically distinct (BA.5.2) SARS-CoV-2 viruses were determined 14 days post second immunization by micro-neutralization assays. ns, non-significant; *, *p* < 0.05; **, *p* < 0.01.

**Supplemental Figure 2.**
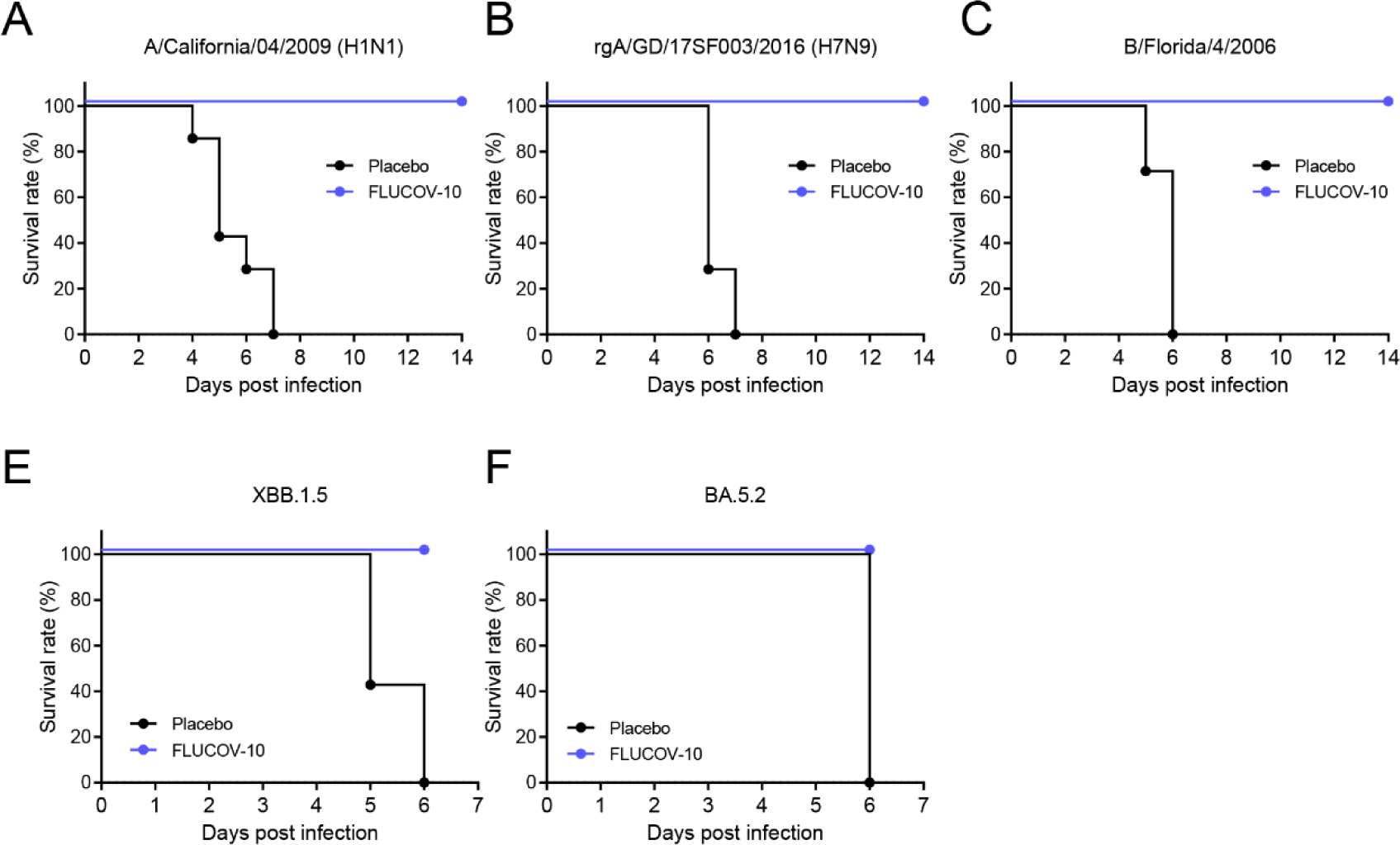
Survival Rates in Mice Immunized with FLUCOV-10 and Challenged with Influenza or SARS-CoV-2 Viruses (related to Figure 4). The mice were challenged 3 weeks post second immunization with indicated viruses and the survival rates were monitored for 7 or 14 days.

## References

1. Organization WH. WHO Coronavirus (COVID-19) Dashboard 2023 [Available from: https://covid19.who.int/.

2. Vasconcelos GL, Pessoa NL, Silva NB, Macedo AMS, Brum AA, Ospina R, et al. Multiple waves of COVID-19: a pathway model approach. Nonlinear Dyn. 2023;111(7):6855–72.

3. 3. Organization WH. Statement on the fifteenth meeting of the IHR (2005) Emergency Committee on the COVID-19 pandemic 2023 [updated 5 May 2023. Available from: https://www.who.int/news/item/05-05-2023-statement-on-the-fifteenth-meeting-of-the-international-health-regulations-(2005)-emergency-committee-regarding-the-coronavirus-disease-(covid-19)-pandemic.

4. Krammer F, Ellebedy AH. Variant-adapted COVID-19 booster vaccines. Science. 2023;382(6667):157-9.

5. Organization WH. Influenza (Seasonal) 2023 [updated 3 October, 2023. Available from: https://www.who.int/news-room/fact-sheets/detail/influenza-(seasonal).

6. Wang Y, Tang CY, Wan XF. Antigenic characterization of influenza and SARS-CoV-2 viruses. Anal Bioanal Chem. 2022;414(9):2841–81.

7. Furlow B. Triple-demic overwhelms paediatric units in US hospitals. Lancet Child Adolesc Health. 2023;7(2):86.

8. Koutsakos M, Kedzierska K, Subbarao K. Immune Responses to Avian Influenza Viruses. Journal of immunology. 2019;202(2):382–91.

9. Organization WH. Ongoing avian influenza outbreaks in animals pose risk to humans 2023 [updated 12 July 2023. Available from: https://www.who.int/news/item/12-07-2023-ongoing-avian-influenza-outbreaks-in-animals-pose-risk-to-humans.

10. Organization WH. Avian Influenza Weekly Update Number 920 2023 [updated 3 November, 2023. Available from: https://cdn.who.int/media/docs/default-source/wpro---documents/emergency/surveillance/avian-influenza/ai_20231103.pdf?sfvrsn=5bc7c406_33.

11. Ma Q, Li R, Guo J, Li M, Ma L, Dai J, et al. Immunization with a Prefusion SARS-CoV-2 Spike Protein Vaccine (RBMRNA-176) Protects against Viral Challenge in Mice and Nonhuman Primates. Vaccines. 2022;10(10).

12. Ma Q, Li M, Ma L, Zhang C, Zhang H, Zhong H, et al. SARS-CoV-2 bivalent mRNA vaccine with broad protection against variants of concern. Front Immunol. 2023;14:1195299.

13. Polack FP, Thomas SJ, Kitchin N, Absalon J, Gurtman A, Lockhart S, et al. Safety and Efficacy of the BNT162b2 mRNA Covid-19 Vaccine. The New England journal of medicine. 2020;383(27):2603–15.

14. Baden LR, El Sahly HM, Essink B, Kotloff K, Frey S, Novak R, et al. Efficacy and Safety of the mRNA-1273 SARS-CoV-2 Vaccine. The New England journal of medicine. 2021;384(5):403–16.

15. Creech CB, Walker SC, Samuels RJ. SARS-CoV-2 Vaccines. Jama. 2021;325(13):1318–20.

16. Karim SSA, Karim QA. Omicron SARS-CoV-2 variant: a new chapter in the COVID-19 pandemic. Lancet. 2021;398(10317):2126–8.

17. Zhao Z, Zhou J, Tian M, Huang M, Liu S, Xie Y, et al. Omicron SARS-CoV-2 mutations stabilize spike up-RBD conformation and lead to a non-RBM-binding monoclonal antibody escape. Nature communications. 2022;13(1):4958.

18. Chalkias S, Feng J, Chen X, Zhou H, Marshall JC, Girard B, et al. Neutralization of Omicron Subvariant BA.2.75 after Bivalent Vaccination. The New England journal of medicine. 2022;387(23):2194-6.

19. Chalkias S, Harper C, Vrbicky K, Walsh SR, Essink B, Brosz A, et al. A Bivalent Omicron-Containing Booster Vaccine against Covid-19. The New England journal of medicine. 2022;387(14):1279–91.

20. Davis-Gardner ME, Lai L, Wali B, Samaha H, Solis D, Lee M, et al. Neutralization against BA.2.75.2, BQ.1.1, and XBB from mRNA Bivalent Booster. N Engl J Med. 2023;388(2):183-5.

21. Zou J, Kurhade C, Patel S, Kitchin N, Tompkins K, Cutler M, et al. Improved Neutralization of Omicron BA.4/5, BA.4.6, BA.2.75.2, BQ.1.1, and XBB.1 with Bivalent BA.4/5 Vaccine. bioRxiv. 2022:2022.11.17.516898.

22. Davis-Gardner ME, Lai L, Wali B, Samaha H, Solis D, Lee M, et al. Neutralization against BA.2.75.2, BQ.1.1, and XBB from mRNA Bivalent Booster. The New England journal of medicine. 2022.

23. Feldman RA, Fuhr R, Smolenov I, Mick Ribeiro A, Panther L, Watson M, et al. mRNA vaccines against H10N8 and H7N9 influenza viruses of pandemic potential are immunogenic and well tolerated in healthy adults in phase 1 randomized clinical trials. Vaccine. 2019;37(25):3326–34.

24. Lee IT, Nachbagauer R, Ensz D, Schwartz H, Carmona L, Schaefers K, et al. Safety and immunogenicity of a phase 1/2 randomized clinical trial of a quadrivalent, mRNA-based seasonal influenza vaccine (mRNA-1010) in healthy adults: interim analysis. Nature communications. 2023;14(1):3631.

25. moderna. Moderna Announces Interim Phase 3 Safety and Immunogenicity Results for mRNA-1010, a Seasonal Influenza Vaccine Candidate 2023 [updated 16 February, 2023. Available from: https://www.accesswire.com/739660/Moderna-Announces-Interim-Phase-3-Safety-and-Immunogenicity-Results-for-mRNA-1010-a-Seasonal-Influenza-Vaccine-Candidate

26. Organization WH. Antigenic and genetic characteristics of zoonotic influenza A viruses and development of candidate vaccine viruses for pandemic preparedness 2022 [updated September 2022. Available from: https://cdn.who.int/media/docs/default-source/influenza/who-influenza-recommendations/vcm-southern-hemisphere-recommendation-2023/202209_zoonotic_vaccinvirusupdate.pdf.

27. Data OWi. SARS-CoV-2 sequences by variant, Nov 20, 2023 2023 [updated November 20, 2023. Available from: https://ourworldindata.org/grapher/covid-variants-bar.

28. Kurhade C, Zou J, Xia H, Liu M, Chang HC, Ren P, et al. Low neutralization of SARS-CoV-2 Omicron BA.2.75.2, BQ.1.1 and XBB.1 by parental mRNA vaccine or a BA.5 bivalent booster. Nature medicine. 2023;29(2):344-7.

29. Horimoto T, Nakayama K, Smeekens SP, Kawaoka Y. Proprotein-processing endoproteases PC6 and furin both activate hemagglutinin of virulent avian influenza viruses. Journal of virology. 1994;68(9):6074–8.

30. Stieneke-Grober A, Vey M, Angliker H, Shaw E, Thomas G, Roberts C, et al. Influenza virus hemagglutinin with multibasic cleavage site is activated by furin, a subtilisin-like endoprotease. EMBO J. 1992;11(7):2407–14.

31. Wang Y, Lv Y, Niu X, Dong J, Feng P, Li Q, et al. L226Q Mutation on Influenza H7N9 Virus Hemagglutinin Increases Receptor-Binding Avidity and Leads to Biased Antigenicity Evaluation. Journal of virology. 2020;94(20).

32. Dong J, Chen P, Wang Y, Lv Y, Xiao J, Li Q, et al. Evaluation of the immune response of a H7N9 candidate vaccine virus derived from the fifth wave A/Guangdong/17SF003/2016. Antiviral research. 2020:104776.

33. Bao L, Deng W, Qi F, Lv Q, Song Z, Liu J, et al. Sequential infection with H1N1 and SARS-CoV-2 aggravated COVID-19 pathogenesis in a mammalian model, and co-vaccination as an effective method of prevention of COVID-19 and influenza. Signal Transduct Target Ther. 2021;6(1):200.

34. Shi R, Zeng J, Xu L, Wang F, Duan X, Wang Y, et al. A combination vaccine against SARS-CoV-2 and H1N1 influenza based on receptor binding domain trimerized by six-helix bundle fusion core. EBioMedicine. 2022;85:104297.

35. Massare MJ, Patel N, Zhou B, Maciejewski S, Flores R, Guebre-Xabier M, et al. Combination Respiratory Vaccine Containing Recombinant SARS-CoV-2 Spike and Quadrivalent Seasonal Influenza Hemagglutinin Nanoparticles with Matrix-M Adjuvant. 2021:2021.05.05.442782.

36. Ye Q, Wu M, Zhou C, Lu X, Huang B, Zhang N, et al. Rational development of a combined mRNA vaccine against COVID-19 and influenza. NPJ vaccines. 2022;7(1):84.

37. Pfizer. Pfizer and BioNTech Announce Positive Topline Data for mRNA-based Combination Vaccine Program Against Influenza and COVID-19 2023 [updated October 26, 2023. Available from: https://www.pfizer.com/news/press-release/press-release-detail/pfizer-and-biontech-announce-positive-topline-data-mrna.

38. Moderna. Moderna Announces Positive Phase 1/2 Data from mRNA-1083, the Company’s Combination Vaccine Against Influenza and COVID-19 2023 [updated October 4, 2023. Available from: https://investors.modernatx.com/news/news-details/2023/Moderna-Announces-Positive-Phase-12-Data-from-mRNA-1083-the-Companys-Combination-Vaccine-Against-Influenza-and-COVID-19/default.aspx.

39. Tafreshi SH. Efficacy, safety, and formulation issues of the combined vaccines. Expert Rev Vaccines. 2020;19(10):949–58.

40. Tsang TK, Cauchemez S, Perera RA, Freeman G, Fang VJ, Ip DK, et al. Association between antibody titers and protection against influenza virus infection within households. 2014;210(5):684–92.

41. Khoury DS, Cromer D, Reynaldi A, Schlub TE, Wheatley AK, Juno JA, et al. Neutralizing antibody levels are highly predictive of immune protection from symptomatic SARS-CoV-2 infection. 2021;27(7):1205-11.

42. Pardi N, Parkhouse K, Kirkpatrick E, McMahon M, Zost SJ, Mui BL, et al. Nucleoside-modified mRNA immunization elicits influenza virus hemagglutinin stalk-specific antibodies. Nature communications. 2018;9(1):3361.

43. Impagliazzo A, Milder F, Kuipers H, Wagner MV, Zhu X, Hoffman RM, et al. A stable trimeric influenza hemagglutinin stem as a broadly protective immunogen. Science. 2015;349(6254):1301-6.

44. Cowling BJ, Perera R, Valkenburg SA, Leung NHL, Iuliano AD, Tam YH, et al. Comparative Immunogenicity of Several Enhanced Influenza Vaccine Options for Older Adults: A Randomized, Controlled Trial. Clinical infectious diseases : an official publication of the Infectious Diseases Society of America. 2020;71(7):1704–14.

45. Reneer ZB, Bergeron HC, Reynolds S, Thornhill-Wadolowski E, Feng L, Bugno M, et al. mRNA Vaccines Encoding Influenza Virus Hemagglutinin (HA) Elicits Immunity in Mice from Influenza A Virus Challenge. 2023:2023.11.15.566914.

46. Arevalo CP, Bolton MJ, Le Sage V, Ye N, Furey C, Muramatsu H, et al. A multivalent nucleoside-modified mRNA vaccine against all known influenza virus subtypes. Science. 2022;378(6622):899-904.

47. Scheaffer SM, Lee D, Whitener B, Ying B, Wu K, Liang CY, et al. Bivalent SARS-CoV-2 mRNA vaccines increase breadth of neutralization and protect against the BA.5 Omicron variant in mice. Nature medicine. 2023;29(1):247-57.

48. Zeng J, Li Y, Jiang L, Luo L, Wang Y, Wang H, et al. Mpox multi-antigen mRNA vaccine candidates by a simplified manufacturing strategy afford efficient protection against lethal orthopoxvirus challenge. Emerging microbes & infections. 2023;12(1):2204151.

49. Menegale F, Manica M, Zardini A, Guzzetta G, Marziano V, d’Andrea V, et al. Evaluation of Waning of SARS-CoV-2 Vaccine-Induced Immunity: A Systematic Review and Meta-analysis. JAMA Netw Open. 2023;6(5):e2310650.

50. Markov PV, Ghafari M, Beer M, Lythgoe K, Simmonds P, Stilianakis NI, et al. The evolution of SARS-CoV-2. Nature reviews Microbiology. 2023;21(6):361–79.

51. Doyon-Plourde P, Przepiorkowski J, Young K, Zhao L, Sinilaite A. Intraseasonal waning immunity of seasonal influenza vaccine - A systematic review and meta-analysis. Vaccine. 2023;41(31):4462–71.

52. Petrova VN, Russell CA. The evolution of seasonal influenza viruses. Nature Reviews Microbiology. 2018;16(1):47-+.

53. Wiemken TL, Khan F, Puzniak L, Yang W, Simmering J, Polgreen P, et al. Seasonal trends in COVID-19 cases, hospitalizations, and mortality in the United States and Europe. Scientific reports. 2023;13(1):3886.

54. Skibinski DA, Baudner BC, Singh M, O’Hagan DT. Combination vaccines. J Glob Infect Dis. 2011;3(1):63–72.

55. Tzenios N, Tazanios ME, Chahine M. Combining Influenza and COVID-19 Booster Vaccination Strategy to Improve Vaccination Uptake Necessary for Managing the Health Pandemic: A Systematic Review and Meta-Analysis. Vaccines. 2022;11(1).

56. Belongia EA, Skowronski DM, McLean HQ, Chambers C, Sundaram ME, De Serres G. Repeated annual influenza vaccination and vaccine effectiveness: review of evidence. Expert Rev Vaccines. 2017;16(7):1–14.

57. Thompson MG, Cowling BJ. How repeated influenza vaccination effects might apply to COVID-19 vaccines. Lancet Respir Med. 2022;10(7):636–8.

58. Pan W, Dong Z, Meng W, Zhang W, Li T, Li C, et al. Improvement of influenza vaccine strain A/Vietnam/1194/2004 (H5N1) growth with the neuraminidase packaging sequence from A/Puerto Rico/8/34. Human vaccines & immunotherapeutics. 2012;8(2):252–9.

59. Thran M, Mukherjee J, Ponisch M, Fiedler K, Thess A, Mui BL, et al. mRNA mediates passive vaccination against infectious agents, toxins, and tumors. EMBO Mol Med. 2017;9(10):1434–47.

60. Thess A, Grund S, Mui BL, Hope MJ, Baumhof P, Fotin-Mleczek M, et al. Sequence-engineered mRNA Without Chemical Nucleoside Modifications Enables an Effective Protein Therapy in Large Animals. Molecular therapy : the journal of the American Society of Gene Therapy. 2015;23(9):1456–64.

61. Holtkamp S, Kreiter S, Selmi A, Simon P, Koslowski M, Huber C, et al. Modification of antigen-encoding RNA increases stability, translational efficacy, and T-cell stimulatory capacity of dendritic cells. Blood. 2006;108(13):4009–17.

62. Kariko K, Muramatsu H, Ludwig J, Weissman D. Generating the optimal mRNA for therapy: HPLC purification eliminates immune activation and improves translation of nucleoside-modified, protein-encoding mRNA. Nucleic acids research. 2011;39(21):e142.

63. Patel S, Ryals RC, Weller KK, Pennesi ME, Sahay G. Lipid nanoparticles for delivery of messenger RNA to the back of the eye. J Control Release. 2019;303:91–100.

64. Organization WH. Manual for the laboratory diagnosis and virological surveillance of influenza 2011 [Available from: https://apps.who.int/iris/bitstream/handle/10665/44518/9789241548090_eng.pdf;jsessionid=AF0FFCECE2F1C0F120B2A8C0B281272F?sequence=1.

65. Wang Y, Lenoch J, Kohler D, DeLiberto TJ, Tang CY, Li T, et al. SARS-CoV-2 Exposure in Norway Rats (Rattus norvegicus) from New York City. mBio. 2023;14(2):e0362122.

66. Hadfield J, Megill C, Bell SM, Huddleston J, Potter B, Callender C, et al. Nextstrain: real-time tracking of pathogen evolution. Bioinformatics. 2018;34(23):4121–3.

67. Katoh K, Misawa K, Kuma K, Miyata T. MAFFT: a novel method for rapid multiple sequence alignment based on fast Fourier transform. Nucleic acids research. 2002;30(14):3059–66.

68. Minh BQ, Schmidt HA, Chernomor O, Schrempf D, Woodhams MD, von Haeseler A, et al. IQ-TREE 2: New Models and Efficient Methods for Phylogenetic Inference in the Genomic Era. Molecular biology and evolution. 2020;37(5):1530–4.

